# GO-CRISPR: a highly controlled workflow to discover gene essentiality in loss-of-function screens

**DOI:** 10.1101/2020.06.04.134841

**Authors:** Pirunthan Perampalam, James I. McDonald, Frederick A. Dick

## Abstract

Genome-wide CRISPR screens are an effective discovery tool for genes that underlie diverse cellular mechanisms that can be scored through cell fitness. Loss-of-function screens are particularly challenging compared to gain-of-function because of the limited dynamic range of decreased sgRNA sequence detection. Here we describe **G**uide-**O**nly control **CRISPR** (GO-CRISPR), an improved loss-of-function screening workflow, and its companion software package, **T**oolset for the **R**anked **A**nalysis of GO-**C**RISPR **S**creens (TRACS). We demonstrate a typical GO-CRISPR workflow in a non-proliferative 3D spheroid model of dormant high grade serous ovarian cancer and demonstrate superior performance to standard screening methods. The unique integration of the pooled sgRNA library quality and guide-only controls allows TRACS to identify novel molecular pathways that were previously unidentified in tumor dormancy. Together, GO-CRISPR and TRACS can robustly improve the discovery of essential genes in challenging biological scenarios.

## INTRODUCTION

Gene editing using CRISPR/Cas9 technology has seen widespread adoption across most biomedical disciplines, including cancer research (Lytle et al., 2019; Shalem et al., 2015). In particular, the ability to multiplex CRISPR gene knockouts on a genome-wide scale has stimulated systematic interrogation of cell biology (Wang et al., 2014). Pooled single guide RNA (sgRNA) libraries are used to create single-gene knockouts in individual cells and selective pressure is applied through culture conditions or drug treatment. Genetic deficiencies that produce resistance or susceptibility are quantitated using sgRNA coding sequences as barcodes to compare gene knockout abundance between the start and end of the experiment (Shalem et al., 2014). CRISPR screens can therefore discover functional roles for genes and pathways not suggested by more traditional hypothesis-driven research.

Gain-of-function genome-wide CRISPR screens can lead to several orders of magnitude change in sgRNA sequence abundance because of resistant cell proliferation, unequivocally identifying resistance genes (Cai et al., 2020; Parnas et al., 2015; Sanson et al., 2018). Conversely, loss-of-function is more challenging to quantitate because complete disappearance of sgRNA sequences for a gene may represent technical failure of the screen design, or its execution (Koike-Yusa et al., 2014). In addition, knockout of an individual gene in the chosen culture condition may not cause lethality with complete penetrance (Thyme et al., 2016). Ultimately, identification of essential genes in loss-of-function screens has relied on prolonged periods of cell proliferation to separate the abundance of bystander sgRNA abundance from true deleterious changes (Shalem et al., 2014). For this reason, CRISPR screens have generally utilized rapidly proliferating 2D cell culture conditions. Scenarios such as the tumor microenvironment, metastasis and tumor dormancy, are better assessed in 3D culture models such as multicellular tumor spheroids or organoids (Fujii et al., 2016; Jacob et al., 2020; Kenny et al., 2015; Vlachogiannis et al., 2018). However, the inability of organoids to quantitatively regenerate from individual cells upon subculture has prevented robust library representation (Ringel et al., 2020), and in some cases this has been compensated by screening more compact, partial genome libraries (Planas-Paz et al., 2019). Furthermore, most 3D spheroids exhibit slower growth kinetics due to hypoxia and necrosis which can further hamper detection of gene loss events (Zanoni et al., 2016). All of these factors likely contribute to stochastic loss of guides which can confound loss-of-function studies since current methods cannot distinguish these Cas9-independent events from bona fide loss-of-function due to gene editing. For these reasons, the classification of gene ‘essentiality’ is highly challenging in 3D culture conditions.

Therefore, there is a need for a screening method that can be adapted for a broad range of complex culture conditions that include low proliferation rates to identify essential genes. This motivated us to develop **G**uide-**O**nly control **CRISPR** (GO-CRISPR). GO-CRISPR is a scalable loss-of-function screening method that can be used to discover essential genes in standard monolayer (2D) or complex 3D culture conditions such as dormant tumor spheroids that exhibit arrested cell proliferation. To support broad usability, we also developed TRACS (**T**oolset for the **R**anked **A**nalysis of GO-**C**RISPR **S**creens) to automate the analysis of GO-CRISPR screens in an easy-to-use software package. Together, GO-CRISPR and TRACS allowed us to discover novel survival pathways in dormant ovarian cancer spheroids, whereas established CRISPR screening and analysis approaches were unable to find essential genes. We expect that this approach can be broadly applied to genome-wide loss-of-function CRISPR screens in low proliferation biological contexts.

## RESULTS

### The GO-CRISPR Workflow

The challenges presented by genome wide CRISPR screening in growth arrested populations of cells motivated us to develop a new workflow that could reveal critical insights into mechanisms of survival in cancer cell dormancy. We developed GO-CRISPR to overcome these challenges and its typical experimental workflow is illustrated in **Figure 1A**. CRISPR screens depend on high-level Cas9 expression to ensure maximum efficiency of gene disruption in Cas9-positive cells transduced with a pooled sgRNA library (L_0_) (Shalem et al., 2014; Zhou et al., 2014). GO-CRISPR uniquely incorporates sequencing data from a parallel screen in which Cas9-negative cells are also transduced with the same pooled sgRNA library (L_0_). Both the Cas9-positive and Cas9-negative cells are treated in an identical manner. Following antibiotic selection for sgRNA transduction and expansion into triplicate cultures, cells are harvested from the initial culture condition (T_0_). Next, both Cas9-positive and Cas9-negative populations are exposed to the desired selective pressure or culture conditions (P_s_) and cells are harvested from the final culture condition (T_f_). Next-generation sequencing (NGS) is then used to quantitate the abundance of PCR-amplified sgRNA sequences from these 12 samples, as well the initial library preparation (L_0_).

**Figure 1:**
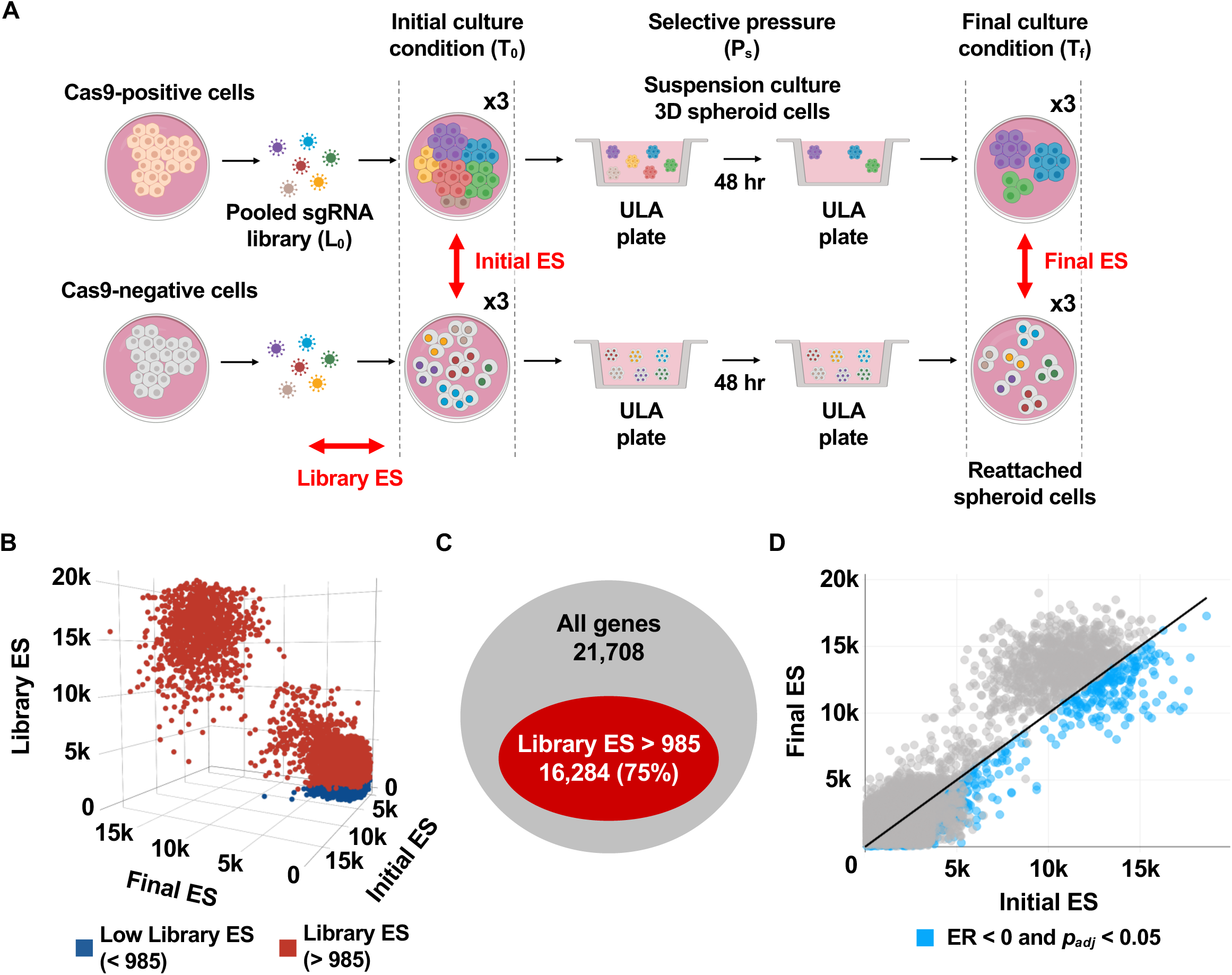
Typical experimental workflow for GO-CRISPR screening and analysis using TRACS. **(A)** iOvCa147 High-grade serous ovarian cancer (HGSOC) cells were transduced with lentivirus expressing Cas9. High efficiency Cas9-positive cells (top row) and Cas9-negative cells (bottom row) were transduced with the GeCKO v2 pooled sgRNA library (L_0_). After antibiotic selection, both Cas9 positive and negative cells were split into triplicates (x3) and maintained in initial culture conditions (T_0_) before being transferred to suspension culture conditions in ULA plasticware (selective pressure, P_s_) to induce spheroid formation and select for cell survival. Viable spheroid cells were then transferred to standard plasticware to facilitate reattachment in the final culture condition (T_f_). The initial pooled sgRNA library (L_0_) and Cas9-positive and Cas9-negative cells were collected at T_0_ and T_f_ for sgRNA quantitation by NGS. TRACS was used to calculate Library, Initial and Final Enrichment Scores (ES) using read quantities from L_0_ and Cas9-positive and Cas9-negative samples. **(B)** 3D plot output from TRACS illustrating the Library ES, Initial ES and Final ES for each gene. Genes highlighted in dark blue have low Library ES (determined by calculating the first quartile value across all Library ES; < 985 in this experiment). **(C)** Euler diagram showing the distribution of retained (in red) and discarded genes based on the Library ES (16,284 genes had Library ES > 985). **(D)** 2D scatter plot output from TRACS showing the distribution of Initial ES and Final ES for all genes. Genes highlighted in light blue (6,717 genes) met the low Library ES cutoff and had a negative Enrichment Ratio (ER) and *p*_*adj*_ < 0.05, indicating their sgRNA abundance decreases in T_f_ compared to T_0_.

To evaluate GO-CRISPR screens, we developed the TRACS algorithm that integrates data from Cas9-positive and Cas9-negative populations to make gene essentiality predictions (**Figure S1**). It is based on assigning gene enrichment scores similar to the single gene score previously described by Wang *et. al*. (Wang et al., 2017). However, TRACS differs by calculating three different enrichment scores for each gene (**Figure 1A** in red). These include a Library Enrichment Score (Library ES) that compares sgRNA read counts for each gene between Cas9-negative cells and the library (L_0_) to determine Cas9-independent non-gene-editing-related changes in abundance. This is an important consideration since pooled sgRNA library preparations do not uniformly represent all genes (Sanjana et al., 2014). An Initial Enrichment Score (Initial ES) is calculated by comparing sgRNA abundances for each gene in Cas9-positive cells relative to their abundances in Cas9-negative cells where they cannot direct gene editing. Lastly, a Final Enrichment Score (Final ES) determines sgRNA abundance between Cas9-positive and Cas9-negative cells following the exposure of both populations to the desired selective pressure or culture condition (P_s_). For each gene, the Library ES, Initial ES and Final ES are weighted according to the number of sgRNAs that are detected for that gene. Thus, a relatively low Initial ES or Final ES indicate reduced sgRNA abundance in the Cas9-positive population and these scores incorporate a penalty for undetected sgRNAs to emphasize the most reliable sgRNA measurements. Finally, TRACS calculates an Enrichment Ratio (ER) that is the log_2_-fold-change (LFC) value between the Final ES and Initial ES to reveal changes in relative abundance between T_0_ and T_f_ culture conditions to detect sgRNAs that were depleted under the selective pressure (P_s_), thereby identifying gene essentiality. The ER informs researchers if a gene shows essentiality for fitness (negative ER) or is non-essential (positive ER) in the experimental condition.

### Discovering Ovarian Cancer Spheroid Vulnerabilities

To demonstrate the value of the GO-CRISPR and TRACS workflow, we performed a genome-wide screen in iOvCa147 high-grade serous ovarian cancer (HGSOC) cells. HGSOC is a highly metastatic disease in which cells detach from primary tumors and aggregate to form 3D spheroids in the abdomen (Matulonis et al., 2016). These spheroid cells are growth arrested and highly resistant to chemotherapy, emphasizing the need to discover their vulnerabilities to improve treatment (Bowtell et al., 2015). We designed a GO-CRISPR screen experiment (**Figure 1A**) to elucidate the genes and pathways that are critical to spheroid cell survival using ultra-low attachment (ULA) plasticware to induce spheroid formation *in vitro* (MacDonald et al., 2017). Ovarian cancer cells undergo significant cell death in suspension culture while spheroids form, therefore after 48-hours we transferred cells back to standard plasticware to allow reattachment and purification of viable cells.

Following analysis with TRACS, we sought to discover genes that were most selectively required for survival in suspension conditions; these represent potential therapeutic targets for dormant ovarian cancer cell spheroids. **Figure 1B** displays each ES in a 3D plot that reveals the distribution of scores in each dimension and highlights genes with low Library ES in dark blue. A low Library ES means that a gene’s sgRNA sequences were poorly represented at T_0_ due to non-gene-editing events that occurred between viral transduction of the pooled sgRNA library and antibiotic selection. This is an important consideration because when a gene’s Library ES is low, its initial sgRNA abundance is also low, and relatively small changes in sgRNA abundance can lead to extreme enrichment scores at T_0_ (Initial ES) or T_f_ (Final ES) (**Figure S2A**). To avoid these false positives, we excluded the first quartile of Library ES from further analysis (Library ES < 985 in this experiment) (**Figure 1C and Figure S2B**). To discover genes essential for spheroid cell survival, we focused our attention on those that had a negative ER. In **Figure 1D**, genes highlighted in light blue met the Library ES cutoff (> 985) and had ER < 0 and *p*_*adj*_ < 0.05 at a false discovery rate (FDR) of 10%. We found 6,717 genes that met these criteria and the top 10 genes with the most negative ER are shown in **Table 1**. This data suggests these are the ten most essential genes required for spheroid cell viability in iOvCa147 cells.

**Table 1:** Top 10 genes with the most negative Enrichment Ratio (ER) in TRACS.

| **Gene** | **Library ES** | **Initial ES** | **Final ES** | **ER** | ***p*<sub>adj</sub>** |
| --- | --- | --- | --- | --- | --- |
| *AGPS* | 2648.39 | 1384.00 | 16.61 | -6.38 | 1.01 x 10<sup>-2</sup> |
| *SLC2A11* | 2421.17 | 2216.28 | 37.33 | -5.89 | 1.06 x 10<sup>-2</sup> |
| *ZC3H7A* | 2397.67 | 2592.22 | 62.67 | -5.37 | 5.19 x 10<sup>-3</sup> |
| *PDCD2L* | 1449.39 | 3018.56 | 82.67 | -5.19 | 1.86 x 10<sup>-3</sup> |
| *NPM1* | 2966.72 | 2070.89 | 68.06 | -4.93 | 9.78 x 10<sup>-3</sup> |
| *KIAA1731* | 3063.72 | 2050.61 | 83.78 | -4.61 | 4.92 x 10<sup>-3</sup> |
| *MAP3K6* | 1741.61 | 2206.22 | 93.39 | -4.56 | 2.76 x 10<sup>-4</sup> |
| *SSH1* | 1281.61 | 1347.67 | 57.50 | -4.55 | 1.64 x 10<sup>-2</sup> |
| *MFN2* | 2779.67 | 2080.00 | 89.83 | -4.53 | 1.16 x 10<sup>-2</sup> |
| *CREBL2* | 1449.78 | 2211.22 | 96.67 | -4.52 | 6.69 x 10<sup>-3</sup> |

### Validation of TRACS Gene Essentiality Predictions

To determine the validity of gene essentiality predictions made by TRACS, we measured its ability to categorize the 1,000 non-targeting control (NTC) sgRNAs from the GeCKO v2 pooled library. These NTC sgRNAs target non-coding intergenic sequences and should rank as non-essential (Sanjana et al., 2014). We computed a receiver operating characteristic curve (ROC) and determined the area under the curve (AUC) was 98.5%, demonstrating that TRACS correctly identified NTC sgRNAs as non-essential (**Figure 2A**). This is a critical control because amplified genome regions produce false essential calls among non-coding controls (Aguirre et al., 2016; Wang et al., 2015). HGSOC is characterized by extensive amplifications and deletions (Patch et al., 2015) and this data demonstrates TRACS eliminates this potentially confounding interpretation. For added validation, we used CRISPR/Cas9 to disrupt the top five genes with the most negative ER (**Table 1)** in iOvCa147 cells. Independent knockout of each gene showed significant loss of viability under suspension culture conditions (**Figure 2B**). Conversely, knockout of the top five genes with the most positive ER did not compromise viability, suggesting our GO-CRISPR screen approach reliably discovers loss-of-function events (**Figure 2C**).

**Figure 2:**
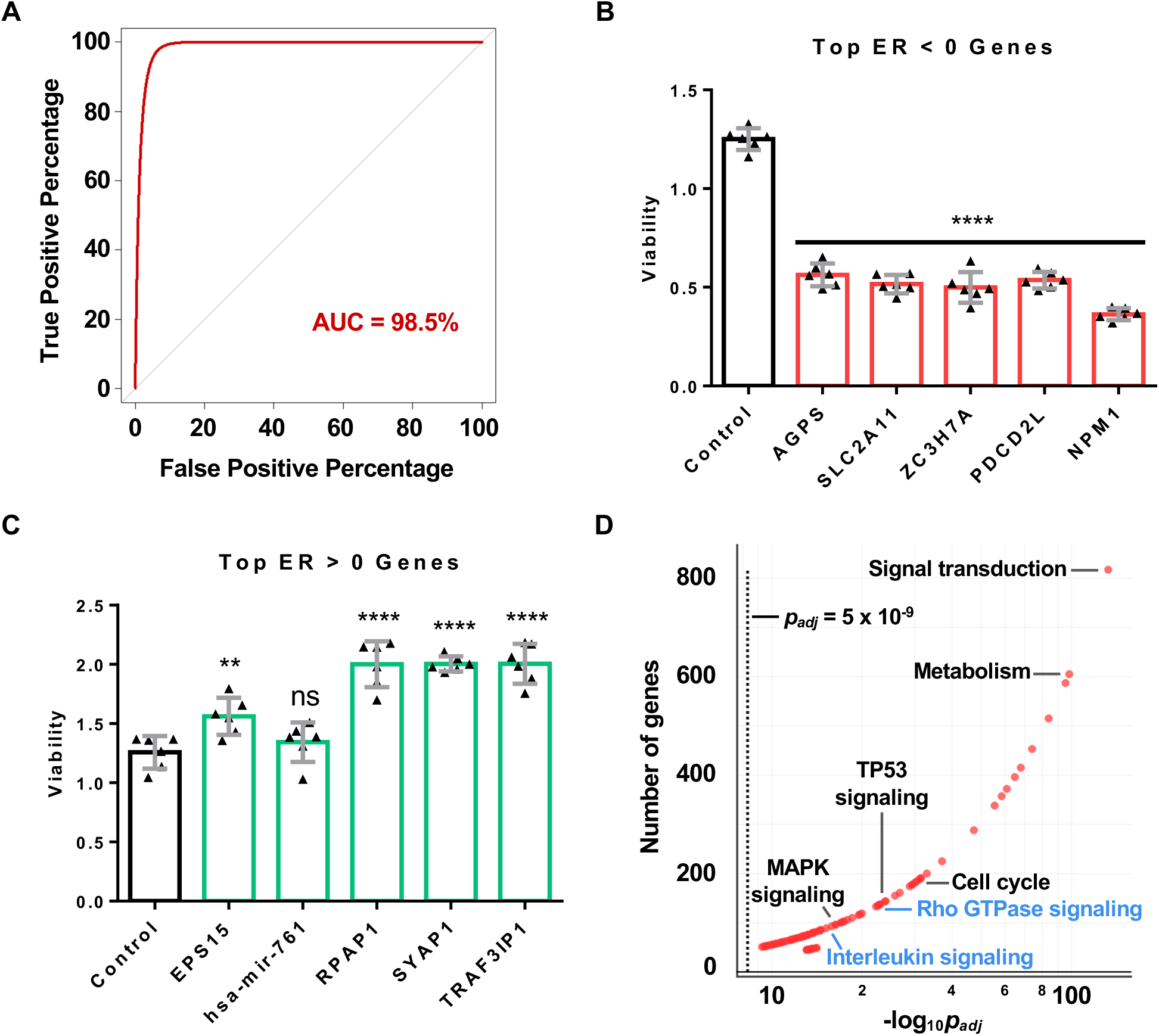
Validation of TRACS gene essentiality predictions. **(A)** The GeCKO v2 pooled library contains 1,000 non-targeting control (NTC) sgRNAs that should not elicit a change in cell fitness. We evaluated the ability of TRACS to classify these sgRNAs by computing a receiver operating characteristic curve (ROC). The area under the curve (AUC) was determined to be 98.5%, indicating TRACS accurately classifies these NTC sgRNAs as non-essential. **(B)** We evaluated the essentiality of the top five genes with the most negative ER: *AGPS, SLC2A11, ZC3H7A, PDCD2L, NPM1* (see **Table 1**). CRISPR/Cas9 was used to disrupt each gene in iOvCa147 cells and pure single-gene knockout populations were assayed for spheroid cell viability in suspension culture conditions. Disruption of these genes resulted in significantly reduced cell viability. Genes in bar graph are arranged from most negative ER to least negative ER. **(C)** We similarly knocked out the top five genes with the most positive ER (*EPS15, hsa-mir-761, RPAP1, SYAP1, TRAF3IP1*) and assayed for viability in suspension culture conditions. Disruption of these genes did not adversely affect viability. Genes in bar graph are arranged from smallest to largest ER. For B and C, each point represents a biological replicate (n = 6). Bars represent means and error bars represent standard deviation. Statistics was performed using one-way ANOVA; ** denotes *p* < 0.01; *** denotes *p* < 0.001; **** denotes *p* < 0.0001; ns denotes not significant (*p* > 0.05). **(D)** We performed gene ontology and pathway enrichment analysis with the 6,717 genes identified by TRACS to have negative ER and plotted the results. The minimum genes required for enrichment per category was set to 45 to ensure stringent selection of pathways. The dashed vertical line represents a *p*_*adj*_ value of 5 × 10^−9^. Pathways to the right of this line have *p*_*adj*_ < 5 × 10^−9^ after controlling for FDR at 10%. Pathways labelled in blue are previously undescribed in HGSOC.

### GO-CRISPR and TRACS Identify Novel Pathways in HGSOC

To further explore the genes identified, we performed gene ontology and pathway enrichment analysis with genes that had a negative ER and *p*_*adj*_ < 0.05 and found 109 significantly enriched pathways (**Figure 2D**). Among these are cell cycle regulation (MacDonald et al., 2017), MAPK signaling (Sun et al., 2017) and TP53 signaling (Patch et al., 2015) which are known to be involved in HGSOC progression and metastasis. Remarkably, our analysis also found novel pathways that have not yet been implicated in HGSOC including Rho GTPase signaling and interleukin signaling. Together, these data demonstrate that GO-CRISPR and TRACS can robustly identify functionally connected genes to enable novel pathway discoveries.

### Comparison of GO-CRISPR with conventional CRISPR screen workflows

Formation of growth arrested HGSOC spheroids in suspension culture is a stressful process in which many cells die without being incorporated into a spheroid. Moreover, the communal nature of spheroids further suggests that individual gene loss events in single cells may be masked in loss-of-function CRISPR screens through non-cell autonomous mechanisms. Thus GO-CRISPR and TRACS were born out of the desire to screen a significantly challenging biological scenario. To fully illustrate the advantages of GO-CRISPR and TRACS, we have analyzed the triplicate replicates of T_0_ and T_f_ solely in Cas9-expressing cells using MAGeCK-RRA, MAGeCK-MLE (Wang et al., 2019) and BAGEL (Hart et al., 2015) as this represents a commonly used CRISPR screen workflow that lacks guide only controls (**Figure S3**). A basic premise for genome-wide CRISPR screens using only Cas9-expressing cells is that poorly represented sgRNAs, or stochastic changes unrelated to the experiment in question, will be removed through statistical cutoffs. Analysis of this data using MAGeCK-RRA/MLE did not detect any essential genes using standard statistical cutoffs (**Figure S3A-D**), including the genes with the most negative ER that were found to be essential by TRACS (**Figure S3I**). BAGEL did not discover essential genes either (**Figure S3G-H**). We then removed statistical cutoffs in MAGeCK-RRA/MLE and found approximately 30% of top-ranked genes had low Library ES according to TRACS, reinforcing the previously described phenomenon of identifying false positives due to low initial sgRNA abundances (**Figure S3E-F and Figure S4**). Additionally, our computed ER discriminates essentiality of NTCs effectively (**Figure 2A**), whereas MAGeCK (without statistical cutoffs) frequently misclassifies NTCs as essential (**Figure S5A**). TRACS was also noticeably more reliable at identifying universally essential and non-essential gene sets (Hart et al., 2015) (**Figure S5B-C**). TRACS penalizes genes that have low sgRNA numbers and favors those with higher sgRNA values to further mitigate the effects of stochastic sgRNA loss and ensure that gene essentiality predictions are made using the largest possible sample size (**Figure S6A**). Without statistical cutoffs, many MAGeCK-MLE top-ranked gene decisions are based on single gRNAs (**Figure S6B**). Overall, integrating data from the pooled sgRNA library and Cas9-negative populations allows TRACS to outperform other methods to accurately predict gene essentiality in a challenging low proliferation, suspension culture scenario.

## DISCUSSION

The GO-CRISPR and TRACS workflow offers an important alternative to conventional genome-wide loss-of-function CRISPR screens. It rigorously identifies genes that contribute to survival and facilitates novel mechanistic discoveries in low proliferation culture conditions by controlling for the stochastic effects of Cas9-independent sgRNA loss. Most notably, we demonstrated the use of this screening workflow in a 3D ovarian cancer spheroid model to identify novel pathways that have not yet been described in HGSOC. This would not have been possible using conventional screening methods that lack guide-only controls.

To accommodate guide-only controls, we needed a new analysis pipeline. CRISPR screen analysis pipelines generally require an understanding of programming or advanced Unix/Linux knowledge to setup and manipulate raw NGS read files. In contrast, the TRACS software suite (https://github.com/developerpiru/TRACS) presents researchers with an easy-to-use graphical environment for analysis and data exploration. TRACS fully automates the analysis process – from raw NGS files to output – which will significantly reduce the barrier for many researchers to use GO-CRISPR. Furthermore, TRACS is fully scalable and can be deployed on a local workstation or a multi-CPU platform such as Amazon Web Services, Google Cloud Platform, or Microsoft Azure. We also provide example workflows and documentation to use TRACS on these platforms, including Docker containers for Linux, Mac OS and Windows that will automate setup.

We used the GeCKO v2 pooled sgRNA library (Sanjana et al., 2014) in our screen. However, the modularity of GO-CRISPR and TRACS will allow for the use of any pooled sgRNA library as long as Cas9 expression is separate from sgRNA viral delivery. In addition, the flexibility of TRACS in terms of unrestricted replicates and sgRNA library size will support the use of validated libraries, such as GeCKO v2, or custom libraries to answer novel questions across biological systems of interest. Taken together, we anticipate GO-CRISPR and TRACS will open new opportunities for loss-of-function screens across diverse model systems and biological questions.

## METHODS

### Generation of Cas9-positive cells

High-grade serous ovarian cancer (HGSOC) iOvCa147 cells have previously been reported (MacDonald et al., 2017). They were transduced with viral particles encoding a Cas9 expression cassette (lentiCas9-Blast, Addgene #52962) to generate cells constitutively expressing Cas9 (Cas9-positive cells). Cells were selected with blasticidin (20 µg/mL). Single-cell clones were isolated by limiting dilution. Lysates were collected from clones and western blots were performed to determine Cas9 expression (Cell Signaling #14697). Cas9 editing efficiency was determined by viability studies using sgRNAs targeting selected fitness genes (*PSMD1, PSMD2, EIF3D*) and a non-targeting control (*LacZ*) as previously reported (Hart et al., 2015). A single clone showing the most effective Cas9 activity was selected for all further studies.

### GeCKO v2 library preparation

HEK293T cells were transfected with the combined A and B components of the GeCKO v2 (Addgene #1000000048, #1000000049) whole genome library (123,411 sgRNAs in total) along with plasmids encoding lentiviral packaging proteins. Media was collected 2-3 days later and any cells or debris were pelleted by centrifugation at 500 x g. Supernatant containing viral particles was filtered through a 0.45 µM filter and stored at −80°C with 1.1 g/100 mL BSA.

### GO-CRISPR screen in iOvCa147 cells

iOvCa147 Cas9-positive or Cas9-negative cells were transduced with virus collected as described above at a multiplicity of infection of 0.3 and with a predicted library coverage of >1000-fold. Cells were grown in media containing 2 µg/mL puromycin (Sigma #P8833) to eliminate non-transduced cells. Cells were maintained in complete media containing puromycin in all following steps. A total of 1.1 × 10^9^ cells were collected and split into three groups consisting of approximately 3.0 × 10^8^ cells each and were cultured for an additional 2-3 days in complete media, then collected and counted. Triplicate samples of 6.2 × 10^7^ cells were saved for sgRNA sequence quantitation at T_0_. The remaining cells (approximately 1.4 × 10^9^/set) were plated at a density of 2.0 × 10^6^ cells/mL in each of twenty 10 cm ULA plates (total of 60 ULA plates). Following 2 days of culture, media containing spheroids was transferred to ten, 15 cm adherent tissue culture plates (total of 30 plates). The next day unattached spheroid cells were collected and re-plated onto additional 15 cm plates. This process was repeated for a total of 5 days at which point very few spheroids remained unattached. The attached cells were collected for DNA extraction and this population represents T_f_. Complete media refers to DMEM/F12 media (Gibco #11320033) supplemented with 10% FBS (Wisent FBS Performance lot #185705), 1% penicillin-streptomycin glutamine (Wisent #450-202-EL) and 2 µg/mL puromycin (Sigma #P8833).

### High-throughput next generation sequencing (NGS)

Cells were harvested and DNA was extracted using QIAmp Blood Maxi Kits (QIAGEN #51194). Genomic encoded sgRNA sequences were PCR amplified as previously described (Joung et al., 2017). Two rounds of PCR were performed. The initial round serves to increase the abundance of the initial sgRNA population, while the second round inserts barcodes necessary for identification of group and replicate number (sample barcode). PCR products were gel purified, quantitated by Qubit (Invitrogen), pooled and sequenced using an Illumina NextSeq 550 75-cycle high output kit (#20024906). FASTQ files were obtained containing raw reads and were demultiplexed to obtain individual FASTQ files for each sample. FASTQ files were processed accordingly for downstream analysis with TRACS, MAGeCK, or BAGEL.

### Analysis with MAGeCK

FASTQ files were trimmed with Cutadapt (1.15) to remove adapter sequences and sample barcode identifiers. The library reference file (CSV) for the GeCKOv2 library was used in Bowtie2 (2.3.4.1) to align the initial library read FASTQ file and generate a BAM file (Samtools 1.7) in order to increase the read depth of the initial library. This library BAM file and trimmed FASTQ files for all samples were then inputted into the MAGeCK (0.5.6) count function to generate read counts. Differences in sgRNA abundance were computed using the MAGeCK-RRA (robust ranking aggregation) or MAGeCK-MLE (maximum likelihood estimation) methods. All plots and comparisons to TRACS were performed in R (3.6.2).

### Analysis with BAGEL

BAGEL (0.91) was run using read counts generated by the MAGeCK (0.5.6) count function as described above. Standard non-essential and essential training gene sets were used as previously described (Hart et al., 2015). Bayes factors (BFs) obtained by BAGEL were plotted in R (3.6.2).

### Analysis with TRACS

The library reference file containing a list of all sgRNAs and their sequences (CSV file), raw reads for the pooled sgRNA library (FASTQ files (L_0_) and raw reads (FASTQ files) for all Initial (T_0_) and Final (T_f_) replicates for Cas9-positive and Cas9-negative cells (12 replicates) were loaded into TRACS (https://github.com/developerpiru/TRACS). TRACS then automatically trimmed the reads using Cutadapt (1.15). TRACS builds a Bowtie2 (2.3.4.1) index and aligns the trimmed initial sgRNA library read file to generate a BAM file using Samtools 1.7. MAGeCK (0.5.6) is then used to generate read counts from this library BAM file and all the trimmed sample FASTQ files. Instead of dropping all reads below a certain threshold (*e*.*g*. <30 counts), all reads were incremented by 1 to prevent zero counts and division by zero errors. The TRACS algorithm was then run using this read count file to determine the Library Enrichment Score (ES), Initial ES, Final ES and the Enrichment Ratio (ER) for each gene (see *The TRACS algorithm* section).

### Data exploration using VisualizeTRACS

The VisualizeTRACS feature, part of the TRACS software suite, was then used to visualize and explore the data output from TRACS. Gene filtering (Library ES > 985, ER < 0, *p*_*adj*_ < 0.05 for our example ovarian cancer workflow) was performed, figures were generated and the final table of essential genes that met these criteria were downloaded for further analysis.

### The TRACS algorithm

After read count preprocessing, TRACS first determines a Gene Score, *GS*, for every gene in the supplied library reference file by calculating the log_2_-fold-change (LFC) from all sgRNAs for that gene for *n* replicates (minimum of 2 replicates required) of Cas9-positive and Cas9-negative samples:

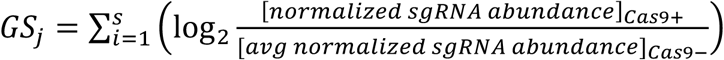

Where *s* is the number of unique sgRNAs for a gene *j*. This is done for each replicate such that for *n* replicates, there are *n* gene scores, *GS*, for a gene *j*. For each *n* replicates, the *GS* for all genes are then ranked in ascending order from 1 to *x*, where *x* is the rank of the gene with the highest *GS* in each respective replicate. TRACS then determines the Enrichment Score, *ES*_*j*_, which is the average rank across all *n* replicates of a gene *j*, divided by the total number of sgRNAs, *s*, identified for that gene.

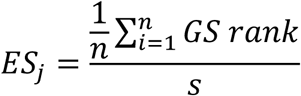

TRACS then determines the Enrichment Ratio, *ER*, for gene *j* by determining the LFC of the 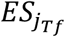 compared to 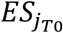.

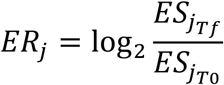

TRACS calculates the *p* value for each gene using a paired t-test by pairing each of the *n* replicates together per gene per the initial (T_0_) condition and final (T_f_) condition. The Benjamini–Hochberg procedure is used to control the false discovery rate (FDR) at the user-defined level (10% in our example workflow).

After the ER is calculated, TRACS determines the distribution of Library ES values across all genes. The cutoff value for the Library ES was set to the first quartile for our example screen.

### Pathway analysis

Using the final list of essential genes from TRACS, we performed gene ontology and pathway enrichment analysis using the ConsensusPathDB enrichment analysis test (Release 34 (15.01.2019)) for top-ranked genes of interest. *p*_*adj*_ values and ER values for each gene were used as inputs. The minimum required genes for enrichment was set to 45 and the FDR-corrected *p*_*adj*_ value cutoff was set to < 0.01. The Reactome pathway dataset was used as the reference. For each identified pathway, ConsensusPathDB provides the number of enriched genes and a *q* value (*p*_*adj*_) for the enrichment. Scatter plots were generated in R (3.6.2) using these values to depict the significant pathways identified.

### Generation of single-gene knockouts

Gibson Assembly (NEB #E2611) was used to clone a pool of four sgRNAs per gene (*AGPS, SLC2A11, ZC3H7A, PDCD2L, NPM1, EPS15, hsa-mir-761, RPAP1, SYAP1, TRAF3IP1*, and *EGFP*) into lentiCRISPR v2 (Addgene #52961). iOvCa147 cells were transduced with viral particles encoding a Cas9 and sgRNA expression cassettes. Cells were selected for 2-3 days in media containing 2 µg/mL puromycin. Knockout cells were cultured for 72 hours in suspension conditions using ULA plasticware (2 × 10^6^ cells per well) to induce spheroid formation. Spheroid cells were then collected and transferred to standard plasticware to facilitate reattachment for 24 hours. Reattached cells were fixed with fixing solution (25% methanol in 1x PBS) for 3 minutes. Fixed cells were incubated for 30 minutes with shaking in staining solution (0.5% crystal violet, 25% methanol in 1x PBS). Plates were carefully immersed in ddH_2_O to remove residual crystal violet. Plates were incubated with detaining solution (10% acetic acid in 1x PBS) for 1 hour with shaking to extract crystal violet from cells. Absorbance of crystal violet at 590 nm was measured using a microplate reader (Perkin Elmer Wallac 1420) for each knockout and normalized to the EGFP control. Percent survival is inferred from relative absorbance.

### Statistics

All error bars in the bar graphs represent standard deviation. Statistical significances were determined using two-way ANOVA. * denotes P < 0.05, *** denotes P < 0.001, **** denotes P < 0.0001 and ns denotes not significant (P > 0.05).

## Abbreviations

AUC: Area under the curve
ER: Enrichment Ratio
ES: Enrichment Score
FDR: False discovery rate
GO-CRISPR: Guide-Only control CRISPR
GUI: Graphical user interface
HGSOC: High-grade serous ovarian cancer
L_0_: pooled sgRNA library
LFC: Log_2_-fold-change
NGS: Next generation sequencing
*p*_*adj*_: adjusted *p* value
P_s_: selective pressure
ROC: Receiver operating characteristic curve
SD: Standard deviation
sgRNA: single guide RNA
TRACS: Toolset for the Ranked Analysis of GO-CRISPR Screens
T_0_: initial culture condition
T_f_: final culture condition
ULA: Ultra-low attachment

## Data and Code Availability

High throughput sequencing data from the GO-CRISPR screen is available from the GEO repository (accession number GSE150246). TRACS is available for download from the GitHub repository at https://github.com/developerpiru/TRACS or on Docker Hub at https://hub.docker.com/r/pirunthan/tracs. Complete documentation, reference sgRNA library file and TRACS output files are also available on the GitHub repository.

## Authors’ contributions

PP wrote the software, analyzed and validated the data and wrote the manuscript. PP and JIM performed the experiments. JIM and FAD wrote the manuscript.

