## Supplemental figures and legends for "GO-CRISPR: a highly controlled workflow to discover gene essentiality in loss-of-function screens"

### Supplementary Figure S1

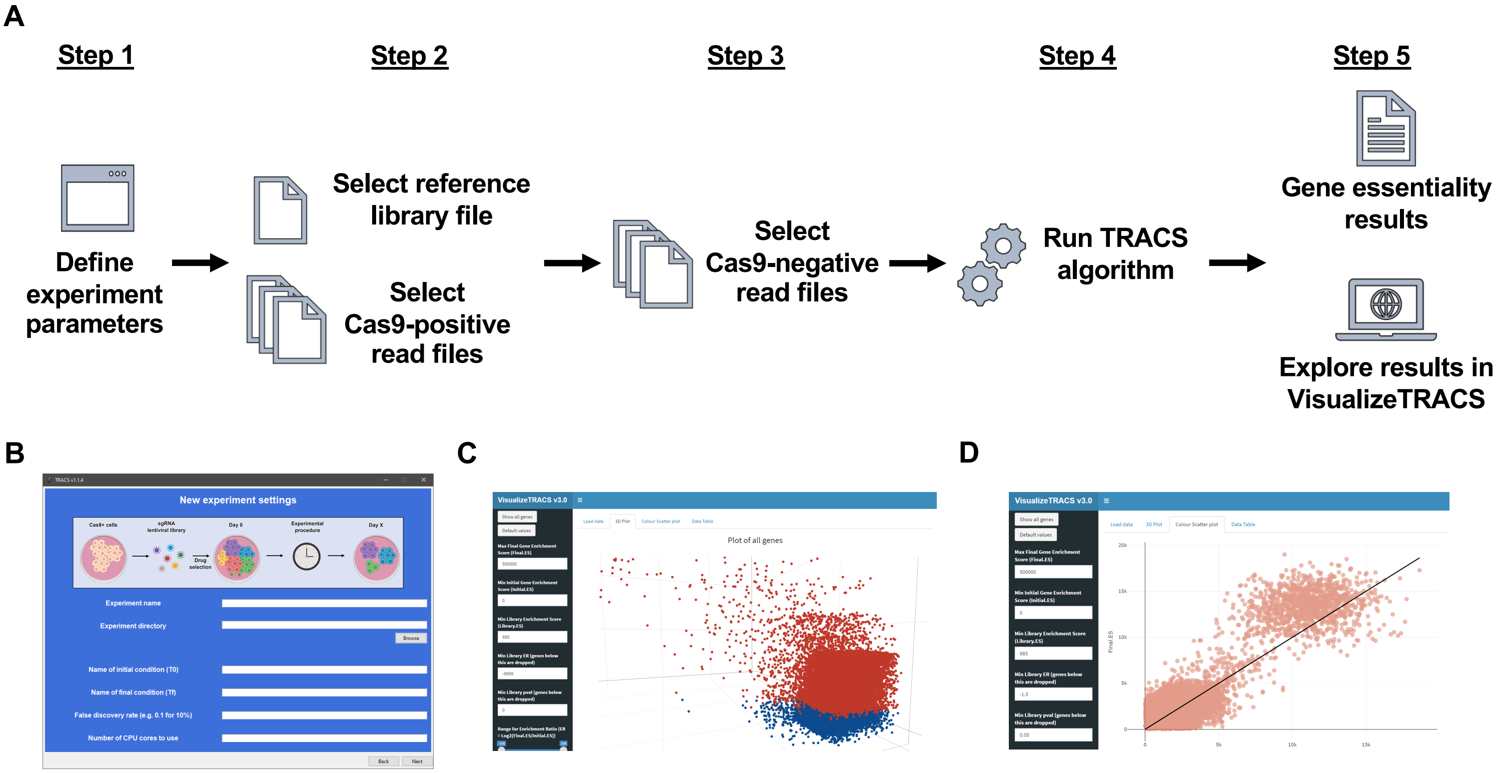

**Supplementary Figure S1: Typical analysis workflow using TRACS to identify essential genes.**

**(A)** The TRACS workflow is separated into five steps. *Step 1:* Experiment parameters are entered in the graphical user interface (GUI). *Step 2:* Library reference file (.csv format) and raw read files (.fastq format) for all Cas9-positive replicates are selected in the GUI. *Step 3:* Raw read files (.fastq format) for all Cas9-negative replicates are selected in the GUI. *Step 4:* Raw reads are trimmed and aligned to generate read counts, then the TRACS algorithm runs to calculate Library ES, Initial ES, Final ES and the ER for each gene. *Step 5:* TRACS saves the results with all scores in an output file which can then be explored using the accompanying VisualizeTRACS data explorer. **(B)** Screenshot of the easy-to-use TRACS GUI asking user to enter experimental parameters (*Step 1*). Subsequent displays provide a similar interface for selecting input data files for *Steps 2-4*. **(C-D)** Screenshot of the accompanying VisualizeTRACS data explorer that researchers can use to visualize and inspect their TRACS output files and generate publication-ready figures. Researchers can control all aspects of filtering and data manipulation (Library ES, Initial ES, Final ES, ER, $p_{adj}$) to fine-tune selection of genes.

Supplementary Figure S2

A

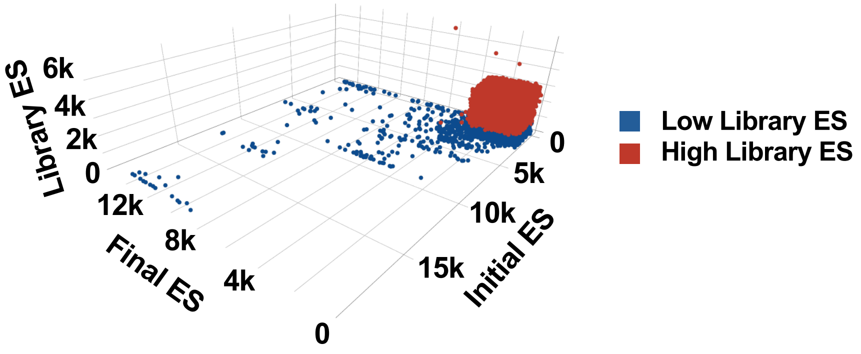


B

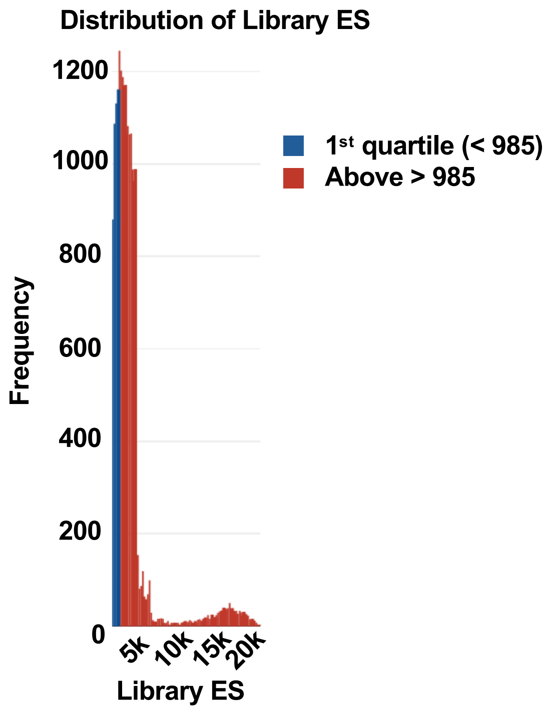

**Supplementary Figure S2: Genes with low Library ES tend to have extreme Initial ES and/or Final ES.**

**(A)** TRACS 3D plot illustrating the distribution of Library ES, Initial ES, Final ES in an extreme case example screen that had very poor representation of sgRNAs at $T_0$ in Cas9-negative cells. Genes that have low Library ES (genes that fall into the first quartile of all Library ES across all genes) are shown in dark blue. These genes also tend to have extreme values for Initial ES and/or Final ES which can lead to potential false positives. This extreme example demonstrates how initial sgRNA abundances can be low due to non-gene-editing events and skew gene scores at $T_0$ (Initial ES) and $T_f$ (Final ES). **(B)** Histogram illustrating the distribution of Library ES across all genes in our GO-CRISPR experiment. To diminish the effects of poorly represented sgRNAs, TRACS determines the distribution of the Library ES across all genes and computes the cutoff value for the first quartile (the bottom 25% of all Library ES; highlighted in dark blue). TRACS then discards genes that have Library ES below this threshold (< 985 for our iOvCa147 screen), however researchers can increase or decrease the threshold within the TRACS software suite for further fine-tuning.

#### Supplementary Figure S3

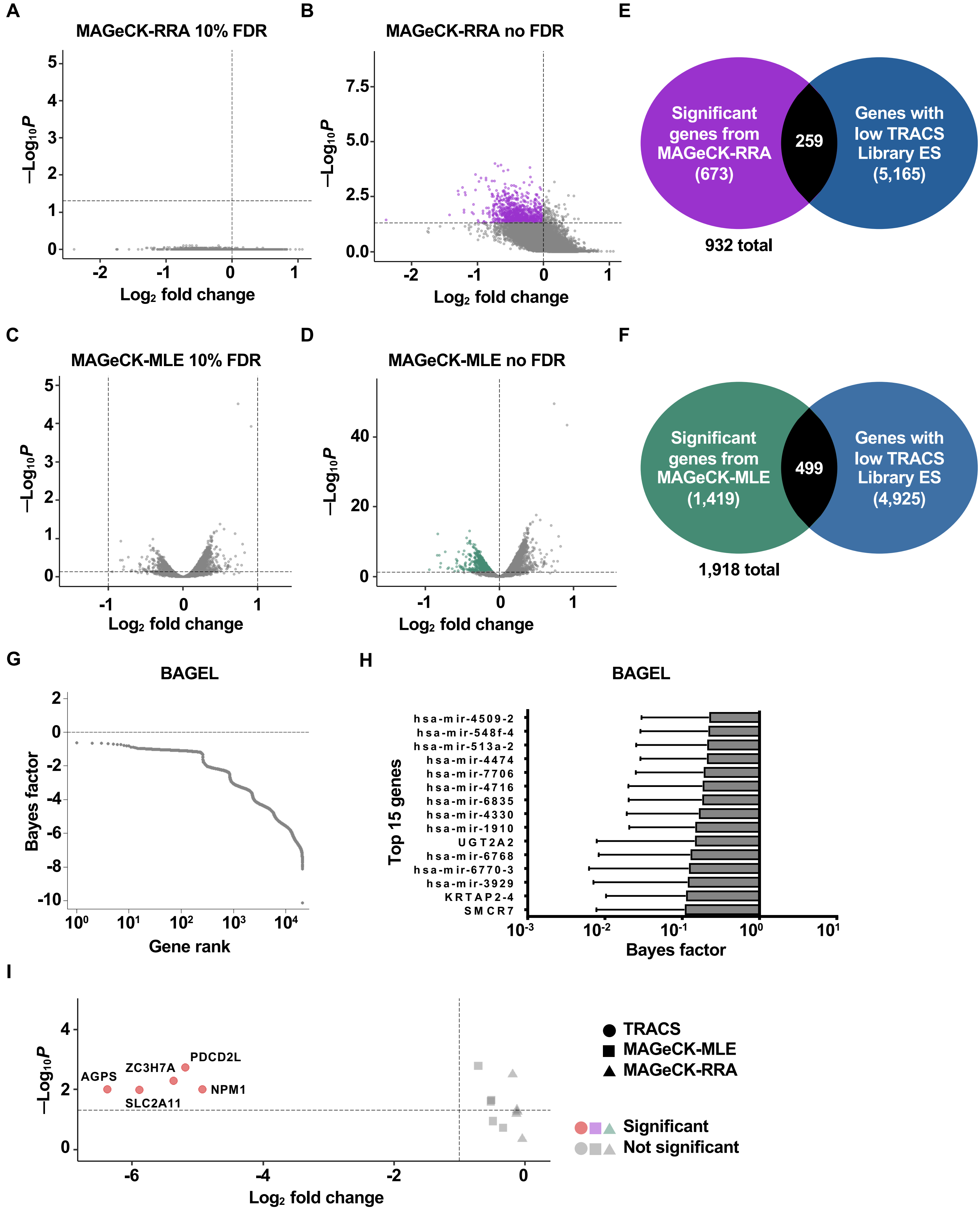

**Supplementary Figure S3: MAGeCK and BAGEL are unable to identify essential genes in our screen using Cas9 positive read data.**

**(A)** We analyzed our screen data using Cas9-positive replicates from $T_0$ and $T_f$ using the MAGeCK-RRA (robust rank aggregation) method with a controlled FDR of 10%. The dashed horizontal line represents $p < 0.05$; any genes above this line are significant. Genes to left of the dashed vertical line have $\log_2$-fold-change (LFC) < 0 indicating their sgRNA abundances decrease from $T_0$ to $T_f$. We did not find any genes to be significant using these typical parameters for MAGeCK-RRA. **(B)** Removal of FDR control with MAGeCK-RRA revealed 932 genes (highlighted in purple) that had LFC < 0 and unadjusted $p$ value < 0.05. Genes shown in grey did not meet these criteria. **(C)** We analyzed our screen data using Cas9-positive replicates from $T_0$ and $T_f$ using the MAGeCK-MLE (maximum likelihood estimation) method with a controlled FDR of 10%. Genes above the dashed horizontal line have $p < 0.05$ and are significant. Genes to the left of the dashed vertical line have LFC < -1, the typically used cutoff for gene essentiality using this method. We did not find any genes that met both of these criteria. **(D)** Removal of FDR control with MAGeCK-MLE and increasing the LFC cutoff to < 0 revealed 1,918 genes (highlighted in green) that had LFC < 0 and $p$ value < 0.05. Genes shown in grey did not meet these criteria. **(E)** Venn diagram showing overlap of the 932 genes (in purple) identified by MAGeCK-RRA with the genes identified by TRACS as having low Library ES (5,424 genes total). 259 genes overlap between the two sets (27.8%). **(F)** Venn diagram showing overlap of the 1,918 genes (in green) identified by MAGeCK-MLE with the genes identified by TRACS as having low Library ES. 499 genes overlap between the two sets (26%). **(G)** We analyzed our screen data using Cas9-positive replicates from $T_0$ and $T_f$ using BAGEL and plotted the Bayes factor output for each gene in relation to the gene ranking. The Bayes factors for all genes were negative, indicating BAGEL did not discover any perturbations in sgRNA abundances between $T_0$ and $T_f$. **(H)** A graphical representation of Bayes factors calculated by BAGEL for the top 15 genes with the highest integer value Bayes factors. All Bayes factors are < 0 indicating gene essentiality was not detected. Error bars show standard deviation for each gene as calculated by BAGEL. **(I)** We explored the output from MAGeCK-RRA and MAGeCK-MLE to determine how each method ranked the top five most essential genes we identified using TRACS. TRACS found these genes to have an ER of at least -4.93 (see Table 1). MAGeCK-RRA and MAGeCK-MLE found these genes to have LFC near 0 and were not significant. The dashed vertical line represents a typical LFC or ER cutoff of -1 and the dashed horizontal line represents a *p* value of 0.05. Genes above and to the left of these lines are significant.

#### Supplementary Figure S4

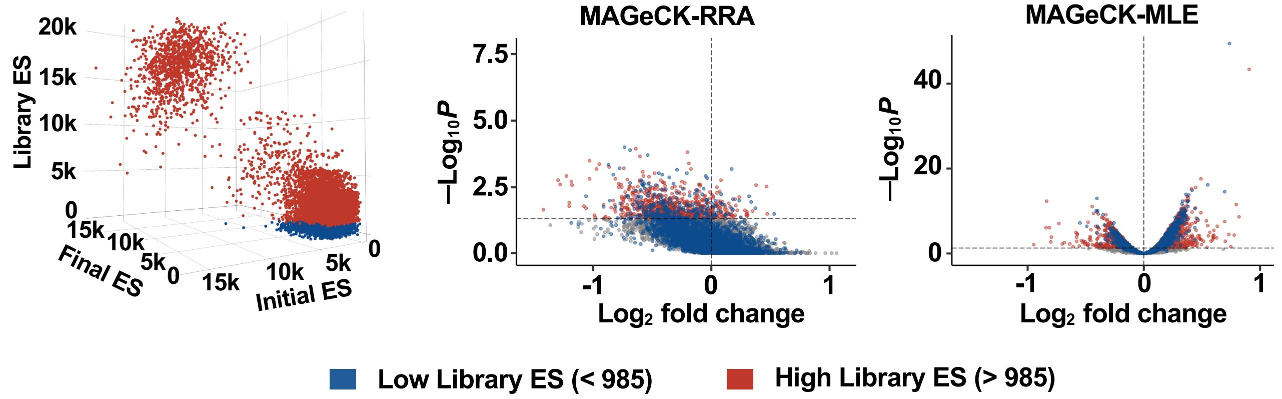

**Supplementary Figure S4: Top-ranked genes by MAGeCK have low representation in the $T_0$ pool of cells.**

The 3D plot highlights in dark blue the genes that TRACS determined to have low Library ES. The vertical axis represents Library ES. The volcano plots illustrate genes that were found to be essential by MAGeCK-RRA or MAGeCK-MLE (LFC < 0 and unadjusted *p* value < 0.05; no FDR cutoffs). Dark blue data points in volcano plots indicate genes that TRACS found to have low Library ES, demonstrating that removing the FDR cutoff selects for genes with poor sgRNA representation. In all three plots, genes in red have Library ES > 985 and genes in dark blue have Library ES < 985.

#### Supplementary Figure S5

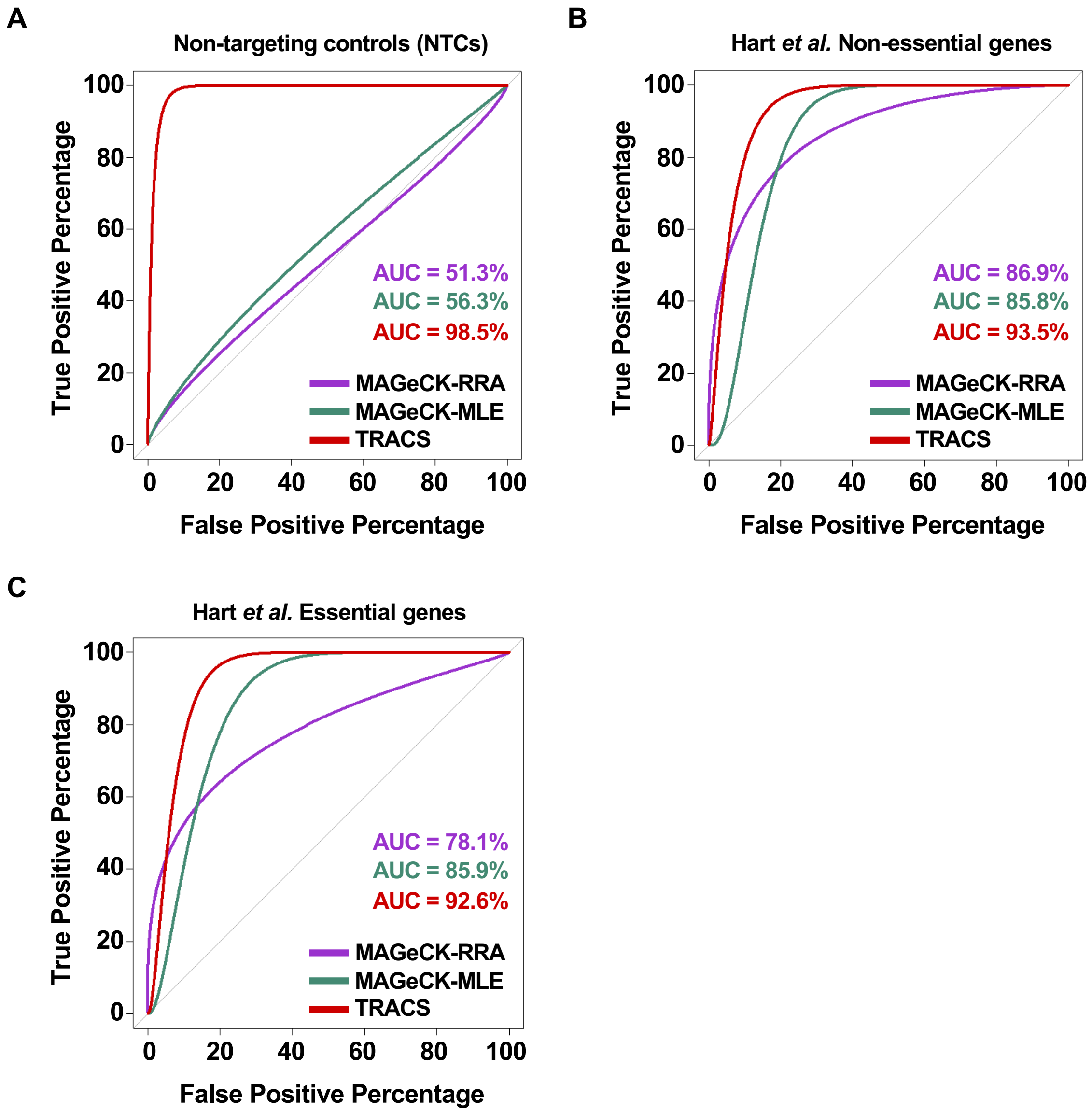

**Supplementary Figure S5: TRACS accurately classifies non-targeting controls and robustly classifies known essential and non-essential gene sets.**

**(A)** We evaluated the ability of MAGeCK to classify the 1,000 NTC sgRNAs in the GeCKO v2 pooled library as non-essential and compared it to TRACS as shown in **Figure 2A**. The AUC for MAGeCK-RRA (51.3%) and MAGeCK-MLE (56.3%) were considerably lower than TRACS (98.5%). **(B)** We evaluated the ability of TRACS and MAGeCK to classify the previously described Hart *et al.* gene set of universally non-essential genes. TRACS (AUC: 93.5%) outperformed MAGeCK-RRA (AUC: 86.9%) and MAGeCK-MLE (AUC: 85.8%) suggesting it can reliably identify these non-essential genes. **(C)** We also evaluated the ability of TRACS and MAGeCK to classify a known set of universally essential genes. TRACS (AUC: 92.6%) consistently outperformed MAGeCK-RRA (AUC:78.1%) and MAGeCK-MLE (AUC: 85.9%) indicating it can robustly identify essential genes.

#### Supplementary Figure S6

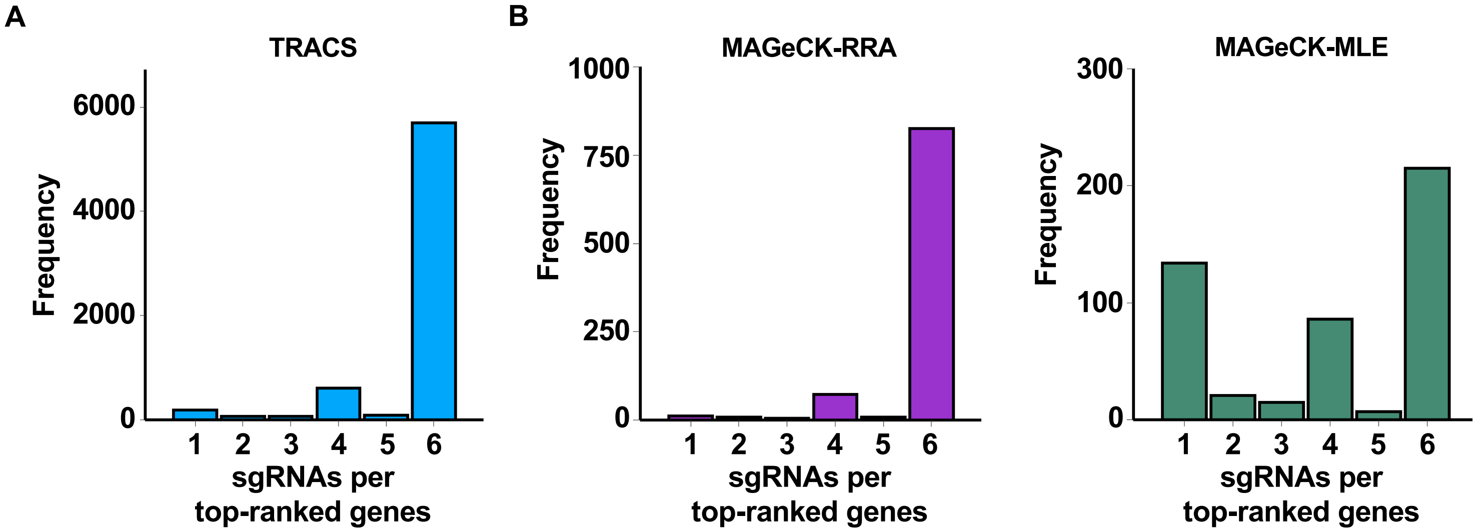

**Supplementary Figure 6: TRACS selects for essential genes based on the most sgRNAs.**

**(A)** Bar plot showing the distribution of the number of sgRNAs per gene for the 6,717 genes that had ER < 0 and $p_{adj} < 0.05$ in TRACS. Light blue color corresponds to light blue data points shown in **Figure 1D**. **(B)** Bar plots showing the distribution of sgRNAs per gene discovered by MAGeCK-RRA and MAGeCK-MLE with LFC < 0 and unadjusted $p$ value < 0.05. Purple and green colors correspond to the colored data points in the volcano plots in **Figure S3**. Most top-ranked genes identified by MAGeCK-RRA had 6 sgRNAs per gene although at reduced frequency which is attributed to fewer genes discovered by MAGeCK. MAGeCK-MLE had wider disparity across genes as it made essentiality calls using as low as 1 sgRNA per gene. The peaks at 4 sgRNAs per gene in each of the three histograms represent miRNAs which have a maximum of 4 sgRNAs instead of 6.
